# Critical role of de novo LTD in the formation and maintenance of hippocampal CA1 place-cell fields

**DOI:** 10.1101/788729

**Authors:** Donovan M Ashby, Jeremy K Seamans, Yu Tian Wang

**Affiliations:** Hotchkiss Brain Institute, University of Calgary; Djavad Mowafaghian Centre for Brain Health, University of British Columbia; Department of Psychiatry, University of British Columbia; Department of Medicine, University of British Columbia

## Abstract

Synaptic plasticity mechanisms may help shape hippocampal place field representations of novel environments, yet direct evidence for how this occurs is lacking. Using multi-channel recordings in freely moving rats, we demonstrate that novelty exploration results in widespread *de novo* long-term depression (LTD) at hippocampal CA1 synapses in a pathway-specific manner, while blockade of LTD expression impairs the maintenance of newly formed place fields. This study therefore reveals an unrecognized role for LTD in the formation and maintenance of hippocampal place fields in novel environments.

## Introduction

The exploration and encoding of a novel environment is a fundamental learning process that occurs on relatively short time scales. It is a useful model for studying how the hippocampus encodes and represents complex, arbitrary associations that are required for episodic memory. Novelty exploration has been demonstrated to promote the induction of electrophysiologically induced long-term depression (LTD) ^1–3^and long-term potentiation (LTP) ^2,4^, the most well-characterized forms of synaptic plasticity in area CA1 of the hippocampus ^5–8^. However, it remains unclear whether novelty exploration can also induce de novo LTP or LTD without exogenous stimulation and if so, what role LTP and/or LTD play as the novel space becomes familiarized and encoded in the hippocampus.

Excitatory pyramidal neurons in areas CA1 and CA3 of the hippocampus consistently show place modulated firing patterns called place fields ^9^. The apparent stability of the place fields upon repeated exposure to the same environment ^10^ suggests that they may have special role in memory. Interventions that affect synaptic plasticity have been shown to also affect the location stability of place cells ^11,12^. However previous studies demonstrating a role for synaptic plasticity in place field maintenance have used interventions that affect other aspects of synaptic transmission or indiscriminately block both LTP and LTD ^12,13^. This leaves open the question about the specific contributions of LTP versus LTD to place field dynamics.

Here, we asked whether novelty exploration would result in *de novo* changes in synaptic strength across multiple recording sites, and if so, were these changes uniform across all sites, as has been reported for a hippocampal dependent aversive learning paradigm ^14^. Synaptic strength was assessed by the response to stimulation by an electrode of the schaffer collaterals. Historically, behaviorally driven changes in the strength of evoked field responses has been confounded by changes in body temperature and behavioral state ^15,16^. To control for these and other unknown variables, we provided an internal control by using a second stimulating electrode to dendritic activate inputs in the striatum oriens. These basal dendritic synapses also exhibit LTP and LTD ^17^, but are not known to be affected by novelty or stress.

## Results

Evoked field potentials at both sites were consistent for one-hour recording sessions across at least 3 days, and were unaffected by brief handling after 30 minutes to simulate environmental transfer (supplementary figure 1). In rats administered saline prior to the recording session, fEPSPs were evoked from stimulation of the stratum oriens (n=37 channels, from 5 rats) and stratum radiatum (n=39 channels, from 5 rats) throughout a 30-minute baseline period in the familiar recording chamber, a subsequent 30-minute period in a novel environment, or a final 30-minute period in the familiar environment (Figure 1). Significant effects of epoch (F(2,148)=41.51, p<.001, group (F(1,74)=22.13, p<.001), and interaction (F(2,148)=3.06, p<.001) were observed. A follow-up comparison between radiatum and oriens evoked potentials during each epoch showed that radiatum evoked fEPSPs exhibited a significant decrease in magnitude relative to oriens stimulation after exposure to the novel environment (84.1%±2.1), a depression that was also maintained during the final 30 minutes in the familiar recording chamber (76.2%±3.0) (Figure 1, t(74)=4.26, p<.001 novel environment, t(74)=4.73 p<.001 familiar environment, vs oriens stimulation). Nearly every channel (38/39 channels) showed a small decrease in amplitude (Figure 1) over the duration of the recording session. On the contrary, oriens evoked potentials showed only a modest, non-significant decrease during novelty exposure (96.9%±2.1) and on return to the familiar environment (95.1%±2.5). These results demonstrate that novel environment induces de novo LTD at the radiatum inputs to the hippocampal CA1 neurons in a pathway specific manner.

**Figure 1.**
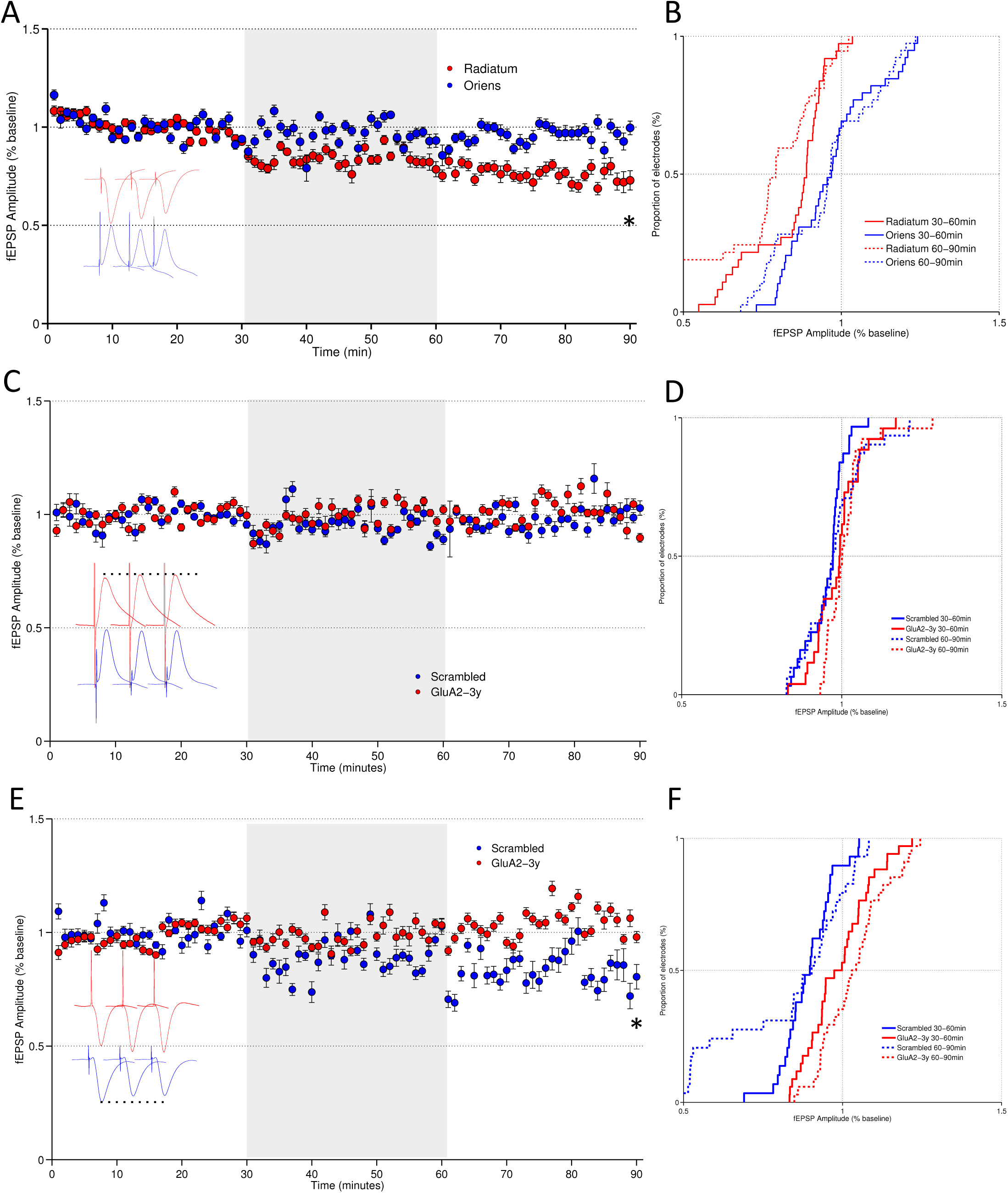
Exposure to a novel environment elicits a decrease in fEPSP specific to stratum radiatum stimulation. (A) A 30-minute baseline recording was followed by 30 minutes in a novel environment (grey shading), followed again by 30 minutes in the familiar recording chamber. A specific decrease in radiatum (red) stimulation was observed with no change in oriens stimulation (blue). (B) Cumulative distribution of amplitude change relative to baseline period for radiatum (red) and oriens (blue) stimulation during novelty (solid) and after returning to the familiar chamber (dashed). In a separate group of animals, tat-GluA2-3y peptide or a scrambled control (2.25 μmol/kg in saline, IV) was delivered 30 minutes prior to baseline recording. (C, D) Stratum oriens stimulation evoked a consistent fEPSP in peptide (red) or scramble (blue) treated rats throughout recording. (E) In radiatum evoked fEPSPs, tat-GluA2-3y (red) blocked de novo LTD observed in the scrambled control group (blue). (F) Similar to saline, scrambled peptide treated rats showed a widespread LTD of radiatum evoked fields after novelty exposure. * indicates p<.05, follow-up comparison between group for epochs, ttest for average fEPSP change.

To test whether the observed *de novo* fEPSP depression was mediated by a facilitation of endocytosis of postsynaptic AMPA receptors at the synapse as the well-characterized LTD induced by electrical stimulation ^18,19^, the GluA2_3Y_ (previously known as GluR2_3Y_) interference peptide or a scrambled control ^18,19^ was administered IV prior to the recording session on the novelty exposure day. Stratum oriens evoked potentials were recorded from five rats administered the GluA2_3Y_ peptide (n=26 channels) and five rats administered the scrambled control peptide (n=31). Stratum radiatum evoked potentials were also recorded from GluA2_3Y_ (n=34 channels) and scrambled treated rats (n=29). Oriens evoked potentials in response to novelty exploration were unaffected by drug treatment (Figure 1), with a modest and non-significant decrease observed during the novel environment in both scrambled (95.5%±1.7) and GluA23y (98.3%±2.9), and no change on return to the novel environment (101.3%±1.5 scrambled, 98.8%±1.1 GluA23y, F(2,110)=3.09, p=.049 main effect of epoch, group and interaction ps>.05). A decrease in evoked potential magnitude was observed in response to radiatum stimulation specifically in scrambled peptide treated rats both during novel environment exploration (88.8%±1.3) and on return to the familiar chamber (83.1%±3.7), consistent with the effect observed with saline treatment (Figure 1). In contrast, no significant change in fEPSP magnitude was seen in rats treated with the active interference peptide either during novel environment exploration (99.8%±1.5) or on return to the familiar chamber (104.8%±1.6, F(2,122)=7.41, p<.001 epoch, F(1,61)=35.99, p<.001 group, F(2,122)=23.91 p<.001 interaction, follow-up comparisons during the novel environment t(61)=5.40, p<.001, and the familiar environment t(61)=5.49 p<.001, GluA2_3Y_ relative to scrambled treatment). Together these results suggest that novel environment exposure produces *de novo* LTD in the stratum radiatum pathway in CA1, that is mediated by activity dependent AMPA receptor endocytosis.

Although some previous studies have reported widespread LTD triggered simply by novel environmental exposure ^1,14^, it was unclear whether this was a specific, synaptically mediated change. We show here that the dominant synaptic change in response to environmental novelty is pathway specific LTD mediated by AMPA receptor endocytosis. The broad nature of this *de novo* LTD suggests that rather than being a direct encoding mechanism for the new environment, this synaptic change may participate in a gain-control type function ^20^ and act in concert with other plasticity changes to encode the new environment. To test this conjecture directly, we next investigated whether blocking LTD during exposure to a novel environment would affect the hippocampal encoding of that environment. Place field formation occurs in a novel environment, and these newly formed place fields are specifically affected by NMDA receptor blockade, CAMKII deletion and protein synthesis inhibition ^12,21–24^. Although these interventions may block both synaptic LTP and LTD, LTP has been suggested to be the mediator of these effects.

Stable place fields are not immediately observable in a novel environment, but form and consolidate over the first several minutes of exploration ^25–28^. The time scale of these dynamics are consistent with the *de novo* LTD we observe. To test potential role of LTD in place field dynamics we first measured place field dynamics a traditional two-dimensional square novel environment (as novel environment exposure induces de nova LTD (Figure 1), and also determined the effects of LTD blockade by GluA2_3Y_ peptide.

A total of 218 separable cells were recorded from area CA1 in eight rats over three recording days in which rats explored a novel box environment on the second day. Place field firing was assessed in a highly familiar environment on one baseline day, and GluA2_3Y_ or a scrambled control was administered prior to recording on the second day, in which a novel environment was presented after a brief exploration of the familiar environment. Following novel environmental exposure, rats were returned to the familiar environment. The same protocol was run on the following day in the absence of any drug administration. In order to examine place field stability across days, matching cells were identified based on waveform characteristics (Figure 2) ^29,30^. Place field stability was measured as a correlation between the firing maps produced in two different sessions from cells identified as matching. A total of 92 cells from eight rats were identifiable across consecutive days in the box environment condition. Within this set, cells were analyzed for correlated place activity across days if they showed a place field within the analyzed environment (peak firing rate exceeding 1 Hz) in at least one recording session. Relatively conserved field firing location was observed in scrambled treated rats in the novel environment (median r=0.58). In contrast, place fields from rats administered the GluA2_3Y_ peptide prior to exposure to a novel environment showed significantly less conservation when compared with fields from the same cells that formed upon re-exploration, although place field conservation was not entirely absent (median r=0.36; t(43)=2.19, p=.034; Figure 2). Treatment with GluA2_3Y_ peptide produced no significant impairment in place field location conservation in the familiar environment as compared with scramble treated controls (scrambled median r=0.56 vs GluA2_3Y_ median r=0.55; t(35)=0.95, p>.05;Figure 2). This indicated that the impaired field location conservation was specific to place fields formed in the novel environment.

**Figure 2.**
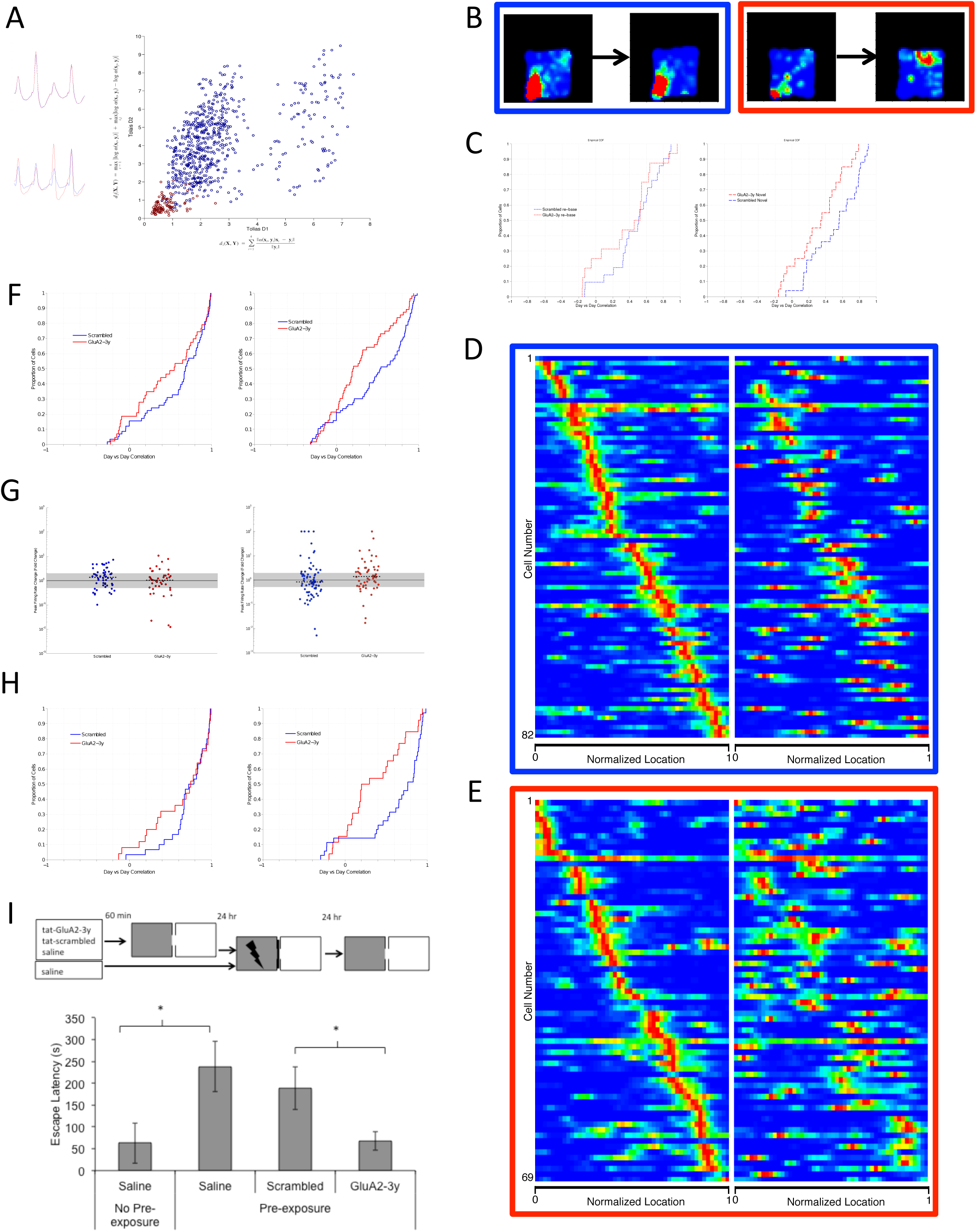
(A) Average waveforms for each sorted cell were compared to average waveforms on subsequent days, recorded from the same tetrode. Projection of computed tolias distance value D1 (x-axis) and D2 (y-axis) indicated a subset of highly similar waveforms (red), classified with a 2-dimensional Bayesian classifier trained on a subset of highly correlated waveforms (p>.97). At left, example tetrode waveforms from non-matched and matched pairs of cells. Each peak represents the average waveform recorded on each of four tetrode channels. (B) Examples of cells recorded from scramble peptide treated (blue) and GluA2-3y peptide treated rats (red), on first exposure to the novel environment and on re-exposure the following day, in a square novel environment. (C) At left, day over day correlation between exposure day and re-exposure day in the familiar recording chamber. Firing fields for cells recorded from scrambled treated rats (blue) and from cells recorded from GluA2-3y (red) treated rats were similarly correlated in the familiar environment p>.05. Drug (2.25 μmol/kg IV) was administered prior to recording only on exposure day between scrambled and GluA23y. At right, day over day correlation between exposure day and re-exposure day in the novel environment. Firing fields for cells recorded from scrambled treated rats (blue) were more highly correlated across days as compared with firing fields from cells recorded from GluA2-3y (red) treated rats. * indicates p<.05 ttest. (D,E) All matched cells recorded in the linear novel novel environment, sorted by firing location on the exposure day. (D) shows cells from scrambled peptide treated rats, with a preserved pattern on the subsequent day. (E) shows cells from GluA2-3y treated rats, with impaired spatial pattern on re-exposure. (F) Day over day correlation in the familiar (left) and novel (right) linear mazes, with peptide (red) and scramble (blue) treatment. GluA2-3y treated rats had significantly reduced correlations relative to scrambled treatment in the novel environment. (G) Peak firing rate change day over day in familiar (left) and novel (right) linear mazes. Median firing rate changes were minimal in all conditions. Grey indicates rate-stable cells, with rate changes between 0.5 and 2. (H) Day over day correlation in rate-stable cells in the familiar (left) and novel (right) linear mazes, with peptide (red) and scramble (blue) treatment. GluA2-3y treated cells were significantly less correlated in the novel environment, even in rate-stable cells. (I) Contextual pre-exposure facilitation of inhibitory avoidance. Top, 60-minutes prior to contextual pre-exposure (8-minutes free exploration of both chambers), rats were administered GluA23y peptide, scrambled control, or saline vehicle (2.25 μmol/kg IV). 24 hours later rats were conditioned by direct placement into the dark compartment and immediate shock (2 × 0.4mA, 0.5s shocks) and removal. 24 hours after training, rats were place in the light compartment and escape latency to the dark compartment was measured. At bottom, rats with no contextual pre-exposure show poor memory relative to pre-exposed rats. GluA23y during pre-exposure impairs inhibitory avoidance conditioning relative to a scrambled control. * indicates p<.05 pairwise comparison following ANOVA.

As place field formation dynamics are more readily measurable in a single dimensional maze where repeated laps of the entire maze are completed more rapidly and with more consistency, we also measured the effect of LTD blockade in a linear maze composed of reconfigurable sections so both extra-maze and intra-maze cues could be manipulated to present multiple different environments (supplementary figure 3). A total of 458 cells from seven rats were recorded during exploration of linear environments. Place field stability in a linear novel environment was assessed using the same criteria as above. Rats administered a scrambled control peptide prior to exposure showed correlated fields between days (median r=0.63, Figure 2). In contrast, place fields from rats administered the GluA2_3Y_ peptide prior to exposure to a novel environment showed significantly less conservation when compared with fields from the same cells that formed upon re-exploration, although place field conservation was again not entirely absent (median r=0.20, t(125)=2.87, p=.008). As observed in the box environment, significantly impaired field conservation due to GluA2_3Y_ peptide treatment was not observed in field recorded in the familiar environment, although a trend was observed (Figure 2; median r=0.73 scrambled, median r=0.54 GluA2-3y; t(97)=1.96, p>.05).

**Figure 3.**
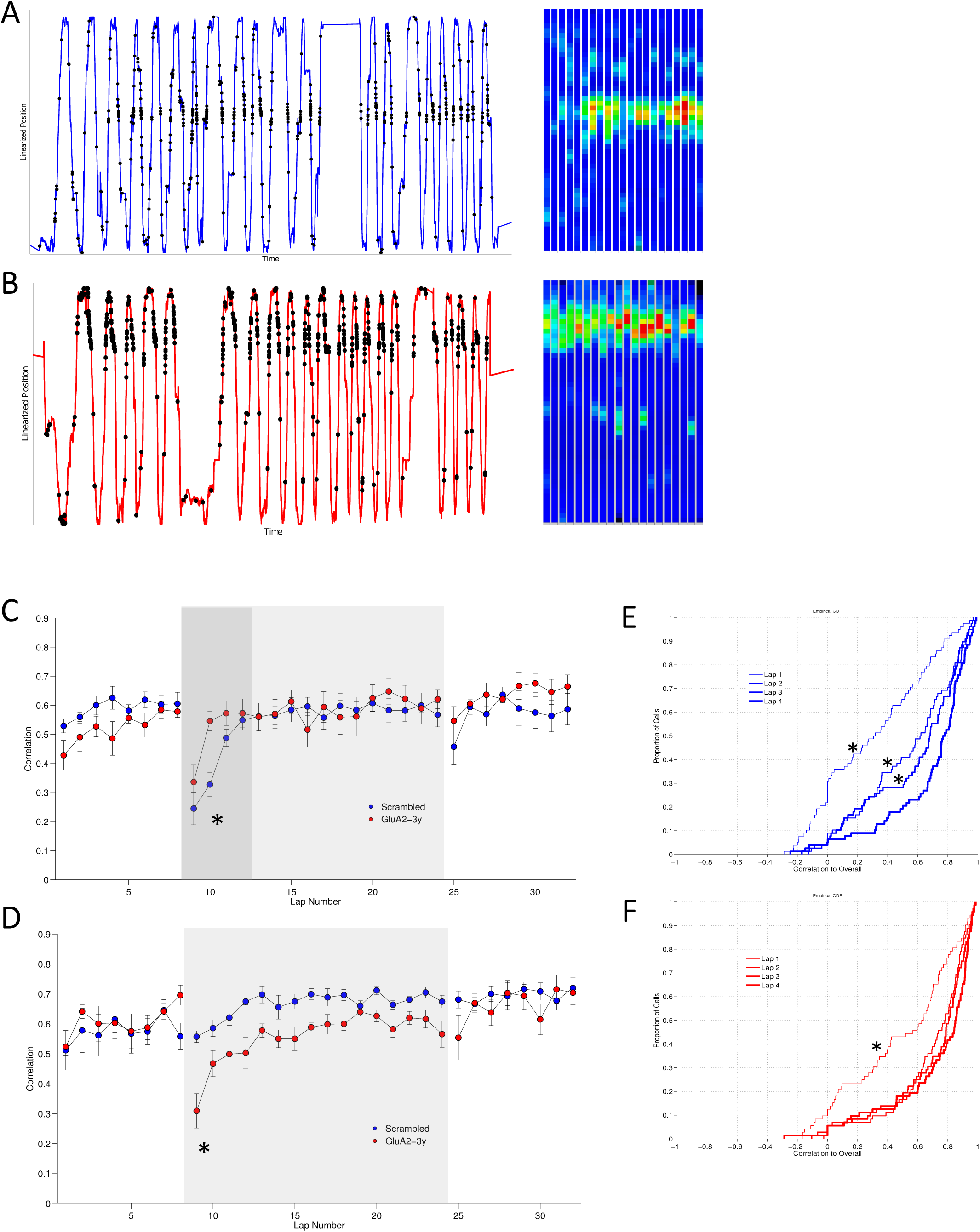
Linearized position and spike locations (left) for one cell from a scramble treated rat (A) and GluA2-3y treated rat (B) and the lap-by-lap firing field (right) upon first exposure to a novel environment. Rats were administered GluA23y or scrambled peptide (2.25 μmol/kg IV) prior to a baseline exploration of the familiar configuration, followed by exploration of a novel configuration (grey box). (C) Place fields were highly correlated and similar between groups in the familiar configuration, but were lower in the first several laps of a novel configuration (dark grey shading). (E,F) The cumulative distribution of correlations on the first four laps were examined. Correlations developed progressively over four trials in scrambled treated rats (E, blue), however highly correlated fields were established after a single lap in GluA23y peptide treated rats (F, red). * indicates p<.05 pairwise comparison only on lap 2 after significant lap by group interaction and p<.05 ks-test vs lap 4. (D) Rats previously administered GluA23y peptide (blue) or scrambled peptide (red) were re-exposed to the novel environment after baseline exploration of the familiar environment. Lap by lap correlations were high in the familiar configuration in both groups, however only rats previously treated with scrambled peptide showed highly correlated fields in the early laps on the previously novel environment (grey shading). In contrast, correlations developed over several laps in rats treated with GluA23y peptide, reminiscent of the initial exposure to the environment in control rats. * indicates p<.05 pairwise comparison after significant group by lap interaction.

Place fields can remap exclusively with a rate change (rate remapping) or by changing field location (global remapping), and reduced correlations in GluA2_3Y_ treated rats could indicate either phenomenon. Rate remapping is normally observed when the same physical locations are superficially altered by changing local cues or environmental shapes. Though exposure and re- exposure to the novel environment in this experiment occurs without any alterations, blocking LTD may preserve the location of place field firing but enhance firing rate changes between sessions. To assess this possibility, firing rate changes between sessions in matched cells were assessed (Figure 2), and the rate-stable subset of cells, where peak firing rates change by less than a factor of 2, were specifically compared. In opposition to the rate-remapping hypothesis, significantly less correlated firing maps were observed in rate-stable cells from peptide treated rats, while firing maps in control rats were highly correlated. Thus, LTD blockade seemed to induce aberrant global remapping of a novel environment.

The effects of LTD blockade on place field stability suggest that LTD blockade should interfere with hippocampal dependent memory, however we have previously shown that the LTD blockade by either GluA2_3Y_ peptide or GluN2B subtype NMDA receptor antagonism does not impair contextual fear conditioning, suggesting that LTD does not play a necessary role in representing contexts ^31^. However contextual fear conditioning is greatly affected by prior habituation to the training context ^32^, and NMDA receptor antagonism in the hippocampus is less effective in blocking inhibitory avoidance in rats subject to context pre-exposure ^33,34^. We sought to address whether blocking LTD during contextual pre-exposure affected the subsequent expression of inhibitory avoidance (Figure 2). Rats that did not receive contextual exposure a day prior to context-shock training (n=8) showed impaired inhibitory avoidance, crossing to the dark chamber within an average of 63 seconds ±46s SEM relative to saline treated rats with contextual pre-exposure (n=8), who showed robust inhibitory avoidance (238±58s, t(14)=2.27, p=.039 vs habituated saline). Scrambled peptide treated rats showed robust inhibitory avoidance (n=14, 188±48 sec), while GluA2_3Y_ treated rats were markedly impaired in inhibitory avoidance (n=15, 68±21 sec, t(27)=2.25, p=.033 vs scrambled). It is important to note that GluA2_3Y_ peptide was administered only during pre-exposure to the environment, leaving LTD unperturbed during context-shock association 24 hours later. Thus, LTD blockade during contextual pre-exposure is sufficient to impair the subsequent context- shock association, supporting a critical role of the novel environment-induced LTD in contextual learning.

As rats covered the entire maze more than once a minute, the linearized maze design permitted an analysis of firing field stability within a session. Previous work shows that stable place fields emerge over the first several laps of a novel linear environment, a timescale consistent with our observations of widespread synaptic LTD. Additionally, experiments that recording field firing in mice exploring virtual environments ^35^ have observed similar dynamics ^36,37^ and implicated dendritic plateau potentials and other nonlinearities in the rapid recruitment of new place fields 38. Importantly, these reports support the idea that an active synaptic plasticity process is required even over the short time span in which this occurs, as the subthreshold voltages recorded before, during, and after recruitment change dramatically. We hypothesized that LTD blockade may interfere with the speed of field recruitment or the early stability of these fields by raising the noise level of synaptic inputs to CA1.

Firing rate maps were computed for each individual lap, and a correlation matrix was calculated for each cell, with the median correlation of each lap to each other lap taken as a similarity index. As rats completed varying numbers of laps within the fixed time recording session, the first eight laps were averaged for familiar configuration sessions (8-minute sessions) and the first 16 laps were averaged for the novel configuration sessions (16-minute sessions). Lap to lap similarly was uniformly high in the familiar environment. Consistent with prior reports, in the novel environment the average place cell firing map increased correlation over the first several laps, with significantly lower correlations vs the final lap until the fourth lap in control rats (Figure 3).

In contrast to our hypothesis, recruitment speed and field stability were not impaired by GluA2_3Y_ peptide treatment, however an alteration in dynamics was observed, with highly correlated place fields were established by the second lap (Figure 3). A comparison of the distribution of correlations on specifically the first four laps demonstrated a rapid, single lap shift to a high correlation distribution in GluA2_3Y_ treated rats (ks-test vs lap 4: lap 1 ks=.37 p<.001, all other laps p>.05), whereas a cumulative shift over four laps was evident in control rats (Figure 3, ks- test vs lap 4: lap 1 ks=.52, p<.001, lap 2 ks=0.29, p=.0017, lap 3 ks=.22 p=.042). This unexpected result indicates that LTD blockade does not diminish the lap to lap stability of place fields, instead seeming to accelerate the formation of a stable place field during novelty exposure. Therefor we report two unique and opposing effects of LTD blockade on the formation of new place field in a novel environment: While stable place fields form more rapidly within the session as compared with controls, place fields are not maintained across days as is typically observed.

## Discussion

These findings illuminate several new features of hippocampal learning and support a role for synaptic long-term depression mechanisms in the acquisition of novel spatial information in the hippocampus. Inhibition of activity-dependent AMPA receptor endocytosis with the peptide inhibitor GluA2_3Y_, a validated and widely used specific inhibitor of long-term depression expression blocked a novelty induced decrease in evoked fEPSPs in the hippocampus, blocked facilitation of inhibitory avoidance by contextual exposure, and impaired the maintenance of place field locations on re-exposure to a novel environment. In addition, acute LTD blockade altered the dynamics of place field formation in a novel environment. Although we demonstrate pathway specific LTD in response to novelty exploration, we blocked LTD at the systemic level and are thus unable to determine whether this specific hippocampal pathway mediates the behavioral and cellular effect we observe, a question that would require pathway specific inhibitors of plasticity to resolve.

There are two important conclusions that can be drawn from the observation that LTD was specifically observed in synapses to the stratum radiatum. The first is that the changes in evoked potential are not likely due to global alterations such as brain state changes or temperature change. The second is that the information transfer via inputs to basal and apical dendrites of CA1 pyramidal cells is differentially altered by novelty. Apical and Basal inputs are physiologically isolated ^39,40^, however both inputs contribute to the generation of the complex spiking in CA1 cells that code for place in a freely moving animal ^41^. It is important to note that apical inputs are collocated with direct inputs from entorhinal cortex, and functional interactions between these inputs influence spiking behavior ^38^ and plasticity in both pathways ^42,43^. LTD in the apical Schaffer collateral input is therefor uniquely positioned to govern the likelihood that coincident inputs from the entorhinal cortex will drive cell spiking ^44^. In the absence of LTD some pre- existing structure may be able to predominate, and we suggest that LTD serves to constrain this structure. This is one possible explanation for the unexpected acceleration of place field formation in rats administered the GluA2-_3Y_ peptide.

Our results are entirely consistent with prior reports of impaired maintenance of place field location after interference with synaptic plasticity ^12^, and suggest that these prior results may be at least partially explained by their effect on LTD, either exclusively or in combination with their effect on LTP. The acquisition of novel spatial information is one of the core functions of the hippocampus and a process to which many major brain systems contribute. A complex cognitive operation is triggered on exposure to a novel environment that begins with novelty recognition and ends with the elaboration of new spatial information. It requires behavioral exploration in concert with hippocampal activity on multiple levels, and we suggest that synaptic LTD serves a specific role in this complex process.

