## Supplementary Information for "Critical role of de novo LTD in the formation and maintenance of hippocampal CA1 place-cell fields"

### Methods

#### Subjects

All experiments were approved by the University of British Columbia Animal Care Committee in accordance with the policies of the Canadian Council on Animal Care. Male sprague-dawley rats (Charles River) weighing 300-450 grams were housed in pairs on a 12hr:12hr standard light cycle, or individually for freely moving electrophysiology experiments. Rats were provided food and water access ad libitum.

#### Drugs

The interference peptide Tat-GluA23y (YGRKKRRQRRR-869YKEGYNVYG877) and scrambled control Tat-Scramble (YGRKKRRQRRR-VYKYGGYNE) were synthesized in house and dissolved in saline (2.25 µmol/kg) for intravenous bolus injection.

#### Microdrive construction

a probe containing 8 bundles of 4 electrodes (25µm tungsten, California Fine Wire) arranged in a 4 by 2 pattern. Within each bundle, electrode tips were cut at 300µm spacing to span the strata of CA1, and bundles were separated by 635µm

Microdrives with four to eight channels were constructed in order to independently adjust the depth of recording tetrodes. A single bundle of 30-gauge stainless steel guide tubes spaced the tetrodes by 300µm. Tetrodes were made of four braided strands of 15µm NiCr wire (California Fine Wire, USA) and sheathed in polyimide tubing (75µm ID). Each actuator of the microdrive consisted of a 12.7mm brass 00-90 hexagonal head machine screw (JI Morris, USA) coupled to a 23-gauge stainless steel drive tube. Tetrodes were threaded through the 30-gauge guide tubes and 23-gauge drive tubes, and individual wires were pinned to a 36 channel electronic interface board (EIB-36N, Neuralynx, USA). Prior to surgery, tetrode tips were 73 precision cut and gold plated with a gold potassium cyanide solution to improve the recording of biopotentials by reducing electrical impedance and tissue reactivity.

#### Surgery

#### Jugular vein catheterization

Rats were anesthetized with isoflurane (5% induction, 2% maintenance) and an incision made in the upper right quadrant of the thorax to expose the right jugular vein. An indwelling silastic catheter (Dow Corning Corp, USA) was inserted into the jugular vein, and the distal end was run subcutaneously to a port on the dorsal surface of the rat between the scapulae. In the case of rats undergoing electrophysiology recordings, the port was integrated into the headcap.

#### Evoked Potential Electrophysiology

Following jugular vein catheterization, anesthetized rats were placed in a stereotaxic frame, and a 3 by 2.5mm cranial window (-1.75 to 4.75mm AP, 47 0.5 to 3.0mm ML) was drilled over the right dorsal hippocampus. Dura was excised to allow implant of a probe... Lateral to the implanted probe, two individually moveable stimulation electrodes (bipolar 50µm stainless steel, AM systems) were implanted into the stratum radiatum and stratum oriens respectively, using standard electrophysiological responses to confirm placement. Electrodes were fixed in place to the skull by dental cement and implant skull screws, which also served as ground and reference points.

CA1 Pyramidal Spiking Electrophysiology

Following jugular vein catheterization, anesthetized rats were placed in a stereotaxic frame, and a 2.0 by 2.0mm cranial window (-2.3 to 4.3mm AP, 1.0 to 3.0mm ML) was drilled over the right dorsal hippocampus. Dura was excised to allow placement of the tip of the microdrive bundle above the cortex overlying the hippocampus. The microdrive was fixed to the skull by dental cement and implant skull screws, which also served as ground and reference points.

Behavioral assays

Evoked field potential recordings

Evoked field recordings in novelty exposure Rats were acclimated to a recording chamber (40x40x60cm) for several days after recovery from surgery. The baseline recording chamber was composed of four uniform dark walls with a removable blue corrugated plastic floor. A single white vertical strip polarized the box for orientation. Extramaze cues were available on all four walls in the recording room. Baseline evoked responses were then tested from 20-200µA stimulation intensity (0.2ms biphasic stimulation), and a stimulation magnitude evoking 50% of the maximal response was used for the remaining recordings. Baseline recordings were made for a minimum of three days (alternating oriens and radiatum test stimulations, 0.016 Hz per site), and the day prior to testing baseline responses were recorded for 1 hour, with brief handling at 30 minutes. On the test day, drugs (saline, tat-GluA23y/tat-Scramble 2.25 µmol/kg) were administered 45 minutes prior to recording. After 30 minutes in the recording chamber, rats were transferred to a novel environment (60x60x60cm polarized box). The novel environment was composed of three black walls and one white wall, with a removable black painted corrugated plastic floor. It was positioned adjacent to the baseline recording box in the same room. Extramaze cues within the room were unchanged. Rats were recorded for 30 minutes in the novel environment, with test stimulation continuing as before, and then transferred back to the baseline recording chamber for 30 minutes. On the following day, rats were re-exposed to the novel environment using the same testing protocol as above, with the exception that no drugs were administered prior to testing.

Inhibitory Avoidance

The inhibitory avoidance apparatus consisted of a light and dark chamber separated by a removable door. Each chamber was 35x30x35cm, and the stainless steel flooring of the dark compartment was connected to a programmable scrambled shock generator (Colbourne Instruments, USA). Inhibitory avoidance was assessed over a three-day protocol designed to isolate the novel exposure to the recording chamber from the chamber-shock association learning (Liang, 1999; Malin & McGaugh, 2006). On day one, rats received a contextual exposure to the chamber, in which they were placed in the light side of the box with the door open and allowed to freely explore both sides of box for 8 minutes. Prior to exploration, rats received bolus IV injection of saline, scrambled peptide, or GluA23y peptide (tat-GluA23y/tatScramble 2.25 µmol/kg). On day two, 24 hours after contextual exposure, rats were placed directly in the dark chamber with the door closed, and received two brief footshocks 2 seconds after placement in the chamber (0.4mA footshock, 0.5 second duration). Five seconds after termination of the footshock rats were removed from the chamber and returned to their homecage. No drugs were administered in the training session. On day three, 24 hours after training, rats were placed in the light compartment facing the far wall, and after 5 seconds the door to the dark chamber was opened. Latency to cross entirely into the dark compartment (all four paws within the compartment) was taken as a measure of inhibitory avoidance learning. A control group of rats was administered saline on day one and trained as above, with the exception that no contextual exposure was performed on day one.

Place Cell recording

In order to record from populations of hippocampal neurons, tetrodes were lowered slowly through the overlying cortex over several weeks of daily sessions while filtered LFP activity (1- 475 Hz) and high frequency unit activity (600-6000 Hz) were monitored. The hippocampal pyramidal layer was identified by the characteristic appearance of ripple activity (150-200 Hz) occurring irregularly during immobility, grooming, and sleep. Tetrodes were slowly lowered 74 further until unit activity was observed during ripples, and subsequent tuning took place to maximize stable, separable units. Place field recording – novel environment During spike tuning, rats were food restricted and trained to forage for sucrose pellets in a recording chamber (40x40x60cm). Baseline recordings were collected during 15-minute foraging sessions to confirm the appearance of place specific firing. On the test day, rats received IV infusions (tat-GluA23y/tat-Scramble 2.25 µmol/kg in 1 ml/kg saline) 45 minutes prior to recording. Rats first ran a 15-minute foraging session in the familiar configuration, then were removed and placed in a novel environment (60x60x60cm polarized box) to run a 30-minute foraging session. Rats were then returned to the baseline recording chamber for an 15-minute session. Place field recording – linear maze Rats were food restricted and trained to shuttle for sucrose pellets delivered to both ends of a linear maze consisting of four 40cm segments (width 12cm, wall height 10cm) linked by turns of 45°, 90°, and 135°. Rats were trained until a minimum of 8 laps were completed within an 8- minute session. On the first exposure day, rats received IV infusions (tat-GluA23y/tat-Scramble 2.25 µmol/kg in 1 ml/kg saline) 45 minutes prior to recording. Rats first ran an 8-minute shuttling session in the familiar maze configuration, then were removed and placed in a novel configuration to run a 16-minute shuttling session. Rats were then returned to the baseline configuration for an 8-minute recording session. 75 On the subsequent re-exposure day, rats ran an 8-minute shuttling session in the familiar maze configuration, then were removed and placed in the same, previously novel configuration to run a 16-minute shuttling session. Rats were then returned to the baseline configuration for an 8-minute recording session. Multiple configurations were possible by adjusting the position of the maze in space and the sequence and direction of turns from one end of the maze to the other.

Data acquisition and analysis

Data acquisition and analysis Local and evoked field potentials were recorded using a 32-channel electrophysiology recording system with integrated headstage preamplifier (Digital Lynx 32, HS-36, Neuralynx, AZ). A native sampling rate of 32kHz was broad-band filtered between 0.1 and 1000 Hz and downsampled to 6.4kHz for analysis. Stimulation (0.2 ms biphasic pulse) was delivered via AM 2100 stimulus isolators, controlled via computer generated TTL pulses. Data was analyzed offline using custom MATLAB analysis code. The amplitude of evoked field potentials was taken as a measure of synaptic strength. As evoked potentials within a recording tract were highly correlated and likely reflected the same population of evoked synapses, only the channel with the highest magnitude potential was used. This also avoided the analysis of complex waveforms recorded from channels intermediate between stratum oriens and stratum radiatum

Place Cell Recording

Spiking and LFPs were recorded using a 32-channel electrophysiology recording system with integrated headstage preamplifier (Digital Lynx 32, HS-36, Neuralynx, AZ), with positional information recorded by tracking a colored LED mounted on the headstage preamplifier with an overhead camera (Neuralynx, AZ). The CA1 pyramidal cell layer was identified using a combination of stereotaxic depth and electrophysiological characteristics. Ripple activity was identified in the LFP channel during quiescence and feeding periods, and tetrode depth was adjusted to maximize multi-unit activity during ripple activity. Spike sorting was conducted offline using manual spike sorting software (Offline Sorter, Plexon). Putative spikes were preprocessed to remove noise and clusters were identified using 3d projections of spike characteristics. Only well-isolated clusters were selected for analysis. Entire recording sessions, whether containing a single or multiple environmental exposures were sorted collapsed over time. Sorted spike data and LFPs were analyzed using an open source MATLAB toolkit (FMAtoolbox) and custom MATLAB analysis code. For each recording session, positional information was first filtered to remove target points occurring outside the arena. Tracking jitters 76 were removed by setting a threshold movement speed. Missing points were interpolated from nearby samples and all position samples were smoothed over a ~2 second Gaussian window. Spikes were velocity filtered to remove firing during immobility, defined as movement below 2.5 cm/s over a smoothed 5-second window. Linearize maze processing For linear track data, two dimensional position samples were linearized to one dimension by collapsing along a single axis defined by a set of vertices corresponding to the end points and three interior corners of the linear maze. Laps were then identified using local extrema on a 25 second smoothed plot of the linearized data. Incomplete laps were removed and subsequent analysis separated inward and outward going laps. Spatial firing rate maps were generated for each isolated unit using a grid of 50x50 bins for 2 dimensional box data, while 1 dimensional linearized data used 50 bins.

Identification of cells across days

The method was based on Tolias et al., (2007) and Powell & Redish (2014). For each cell on each day, the average waveform was calculated, and highly correlated waveforms (r>.97, taken as a 128x1 array of voltage samples, with 32 samples for each tetrode sequentially) recorded from the same tetrode across days in the same rat were taken as a training set of putative matched pairs. Two measures of waveform shape similarity (tolias distance) were calculated for each pair of cells from the same rat, with waveforms expressed as a 32x4 array of voltage samples, each column representing each channel of the tetrode. The first (D1) was a measure of shape similarity of waveforms normalized to minimize the sum of squares difference between the two waveforms on each channel. These scaling factors were used to calculate a normalized Euclidean distance between the waveforms on each channel, summing across channels to produce a single D1 value. A separate D2 value, meant to capture both differences between waveform sizes overall and differences between the scaling factor on each channel was also calculated. Plotted on two dimensions, the D1 and D2 values are both small for matching cell pairs, and larger for non-matching pairs. Using the training set, a Bayesian classifier was used to classify all cell pairs as matching or non-matching. A small minority of cells were positively classified to multiple cell on other days, in this case a linearized tolias distance was used to select only the most closely matched cell pair.

Firing rate maps for pairs of cells were correlated as two linear arrays using pearson's product-moment correlation.

Supplementary Figure Captions

Supplementary Figure 1. Final baseline day evoked field potentials are stable. Two thirty-minute recordings in a familiar recording chamber were separated by a brief handling episode (arrow). Hippocampal CA1 field potentials evoked from stimulation of the oriens (blue) or radiatum (red) pathway showed no significant change across time (A). Inset shows the distribution of evoked field potential change in the second 30-minute epoch relative to the first. Oriens pathway stimulation in groups assigned to scramble (blue) or GluA2-3y peptide treatment (red) showed no change across time during the same recording session on the final baseline day (B). Radiatum pathway stimulation in groups assigned to scramble (blue) or GluA2-3y peptide treatment (red) showed no change across time during the same recording session on the final baseline day (C). It is important to note that no drugs were administered on the baseline days.

Supplementary Figure 2. Re-exposure day evoked field potentials do not show evidence of de novo LTD. A 30-minute recording session in the familiar baseline environment was followed by a 30-minute re-exploration of the novel environment, first explored on the previous day (shaded area). This was followed by a return to the familiar environment for 30-minutes. Stratum oriens evoked field potentials were stable in both scramble treated (blue) and GluA2-3y peptide treated (red) animals (A,B). Stratum radiatum evoked field potentials were similarly stable with average evoked field potentials unchanged during or after re-exploration of the novel environment (C). The distribution of field potentials was significantly different between 30-60 minutes and 60-90 minutes in the scramble treated groups (D, ks-test). * indicates $p<.05$.

Supplementary Figure 3. Place field firing on novel reconfigurations of a linear maze. A familiar configuration of four maze sections was recorded on a baseline day (A). A novel reconfiguration of the maze and external landmarks was presented first on an exposure day, and the same reconfiguration was presented on a re-exposure day. An example cell is shown that fired exclusively in the novel configuration (B). Projecting animal location onto one dimension (C) illustrates the progressive acquisition of a firing field in this cell over several initial laps in the novel maze configuration.

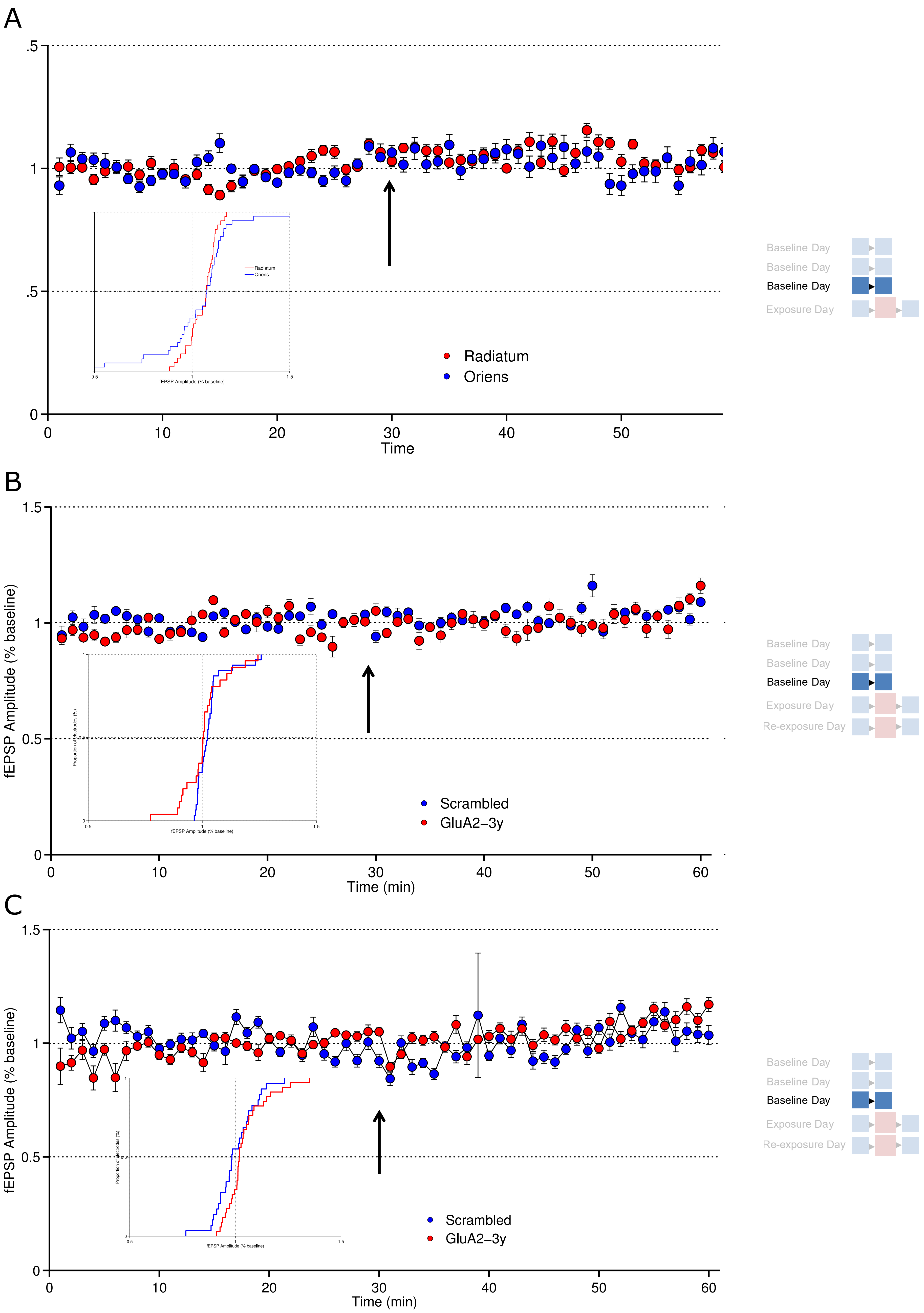
A
.5
1
.5
0
0
10
20
30
40
50
Time
Radiatum
Oriens
fEPSP Amplitude (% baseline)
Baseline Day
Baseline Day
Baseline Day
Exposure Day
B
1.5
1
0.5
0
fEPSP Amplitude (% baseline)
Time (min)
Scrambled
GluA2−3y
Proportion of electrodes (%)
Baseline Day
Baseline Day
Baseline Day
Exposure Day
Re-exposure Day
C
1.5
1
0.5
0
fEPSP Amplitude (% baseline)
0
10
20
30
40
50
60
Time (min)
Scrambled
GluA2−3y
Baseline Day
Baseline Day
Baseline Day
Exposure Day
Re-exposure Day

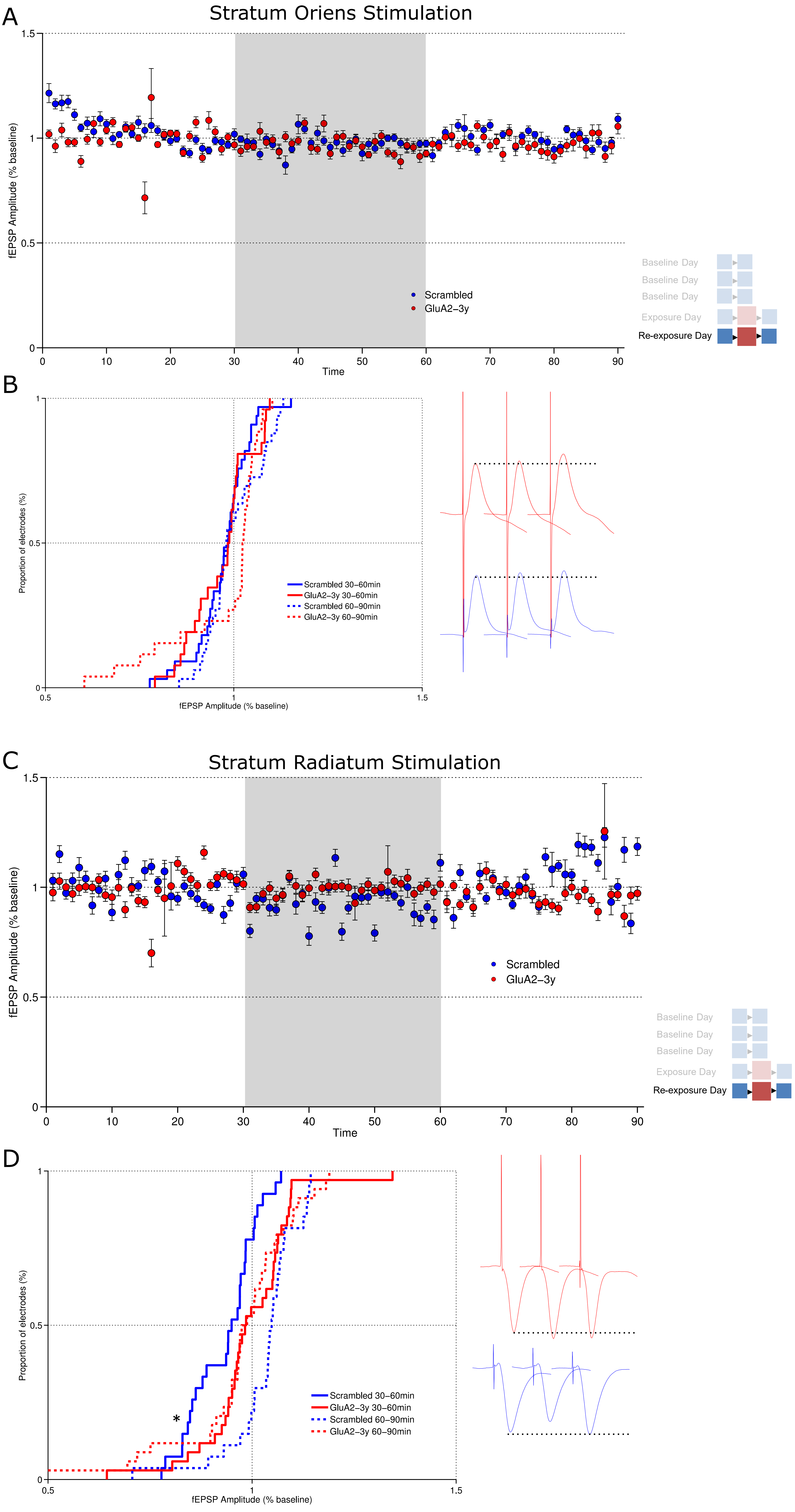
A
Stratum Oriens Stimulation
fEPSP Amplitude (% baseline)
Time
Scrambled
GluA2-3y
Baseline Day
Baseline Day
Baseline Day
Exposure Day
Re-exposure Day
B
Proportion of electrodes (%)
fEPSP Amplitude (% baseline)
Scrambled 30–60min
GluA2-3y 30–60min
Scrambled 60–90min
GluA2-3y 60–90min
C
Stratum Radiatum Stimulation
fEPSP Amplitude (% baseline)
Time
Scrambled
GluA2-3y
Baseline Day
Baseline Day
Baseline Day
Exposure Day
Re-exposure Day
D
Proportion of electrodes (%)
fEPSP Amplitude (% baseline)
Scrambled 30–60min
GluA2-3y 30–60min
Scrambled 60–90min
GluA2-3y 60–90min

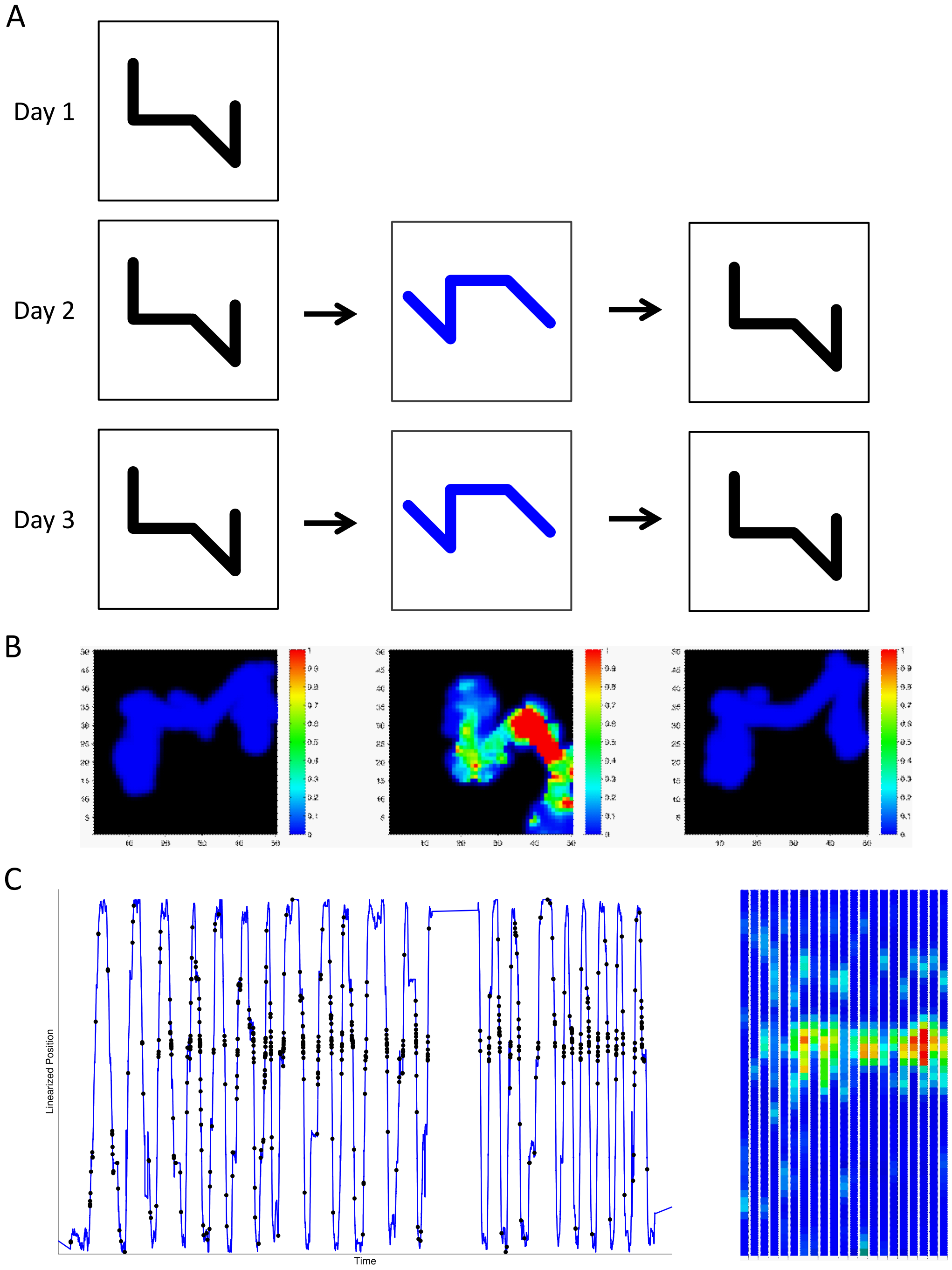
A
Day 1
Day 2
Day 3
B
C
Linearized Position
Time
